# IκBα controls dormancy induction in Hematopoietic stem cell development via retinoic acid

**DOI:** 10.1101/2022.11.17.516971

**Authors:** Roshana Thambyrajah, Zaki Fadlullah, Martin Proffitt, Wen Hao Neo, Yolanda Guillén, Marta Casado-Pelaez, Patricia Herrero-Molinero, Carla Brujas, Noemi Castelluccio, Jessica González, Arnau Iglesias, Laura Marruecos, Cristina Ruiz-Herguido, Manel Esteller, Elisabetta Mereu, Georges Lacaud, Lluis Espinosa, Anna Bigas

## Abstract

Recent findings are challenging the classical hematopoietic model in which long-term hematopoietic stem cells (LT-HSC) are the base of the hematopoietic system. Clonal dynamics analysis of the hematopoietic system indicate that LT-HSC are not the main contributors of normal hemapoiesis in physiological conditions and the hematopoietic system is mainly maintained by multipotent progenitors (MPPs, hereafter HPC) and LT-HSCs are mostly in a non-active state. The first HSCs emerge from the aorta-gonad and mesonephros (AGM) region along with hematopoietic progenitors (HPC) within hematopoietic clusters. Molecular pathways that determine the HSC fate instead of HPC are still unknown, although inflammatory signaling, including NF-κB has been implicated in the development of HSCs. Here, we identify a chromatin binding function for IκBα (also known as the inhibitor of NF-κB) that is Polycomb repression complex 2 (PRC2)-dependent and specifically determines dormant vs proliferating HSCs from the onset of their emergence in the AGM. We find a specific reduction of LT-HSCs in the IκBα knockout new-born pups. This defect is manifested at the FL stage already, and traceable to the first emerging HSCs in the E11.5 AGM, without affecting the general HPC population. IκBα deficient LT-HSCs express dormancy signature genes, are less proliferative and can robustly respond to activation stimuli such as *in vitro* culture and serial transplantation. At the molecular level, we find decreased PRC2-dependent H3K27me3 at the promoters of several retinoic acid signaling elements in the IκBα - deficient aortic endothelium and E14.5 FL LT-HSCs. Additionally, IκBα binding itself is found in the promoters of retinoic acid receptors rarα in the AGM, and rarγ in the LT-HSC of FL. Overall, we demonstrate that the retinoic acid pathway is over-activated in the hematopoietic clusters of IκBα-deficient AGMs leading to premature dormancy of LT-HSCs that persists in the FL LT-HSCs.

## Introduction

HSCs can replenish the adult blood system by generating mature blood cells of all lineages through intermediate stages of multipotent progenitors (Challen et al., 2009). Understanding the molecular programming underlying formation of HSCs during development is critical to improve the feasibility of generating them through Pluripotent Stem cells, or through re-programming of somatic cells to treat blood malignancies (Slukvin II, 2016). Blood stem and progenitor cell (HSPC) emerge through trans-differentiation from specialized endothelial-like cells, termed hemogenic endothelium (HE) and the first cells capable to regenerate the hematopoietic system are found intra-embryonically in the region of the AGM between E10.25 and E11.5 in the mouse embryo (Cumano et al., 2001; Medvinsky and Dzierzak, 1996; North et al., 2002). Here, they accumulate as intra-aortic hematopoietic clusters (IAHC), which are closely associated with the ventral wall of the dorsal aorta (vDA, DA)(de Bruijn et al., 2000, 2002; Jaffredo et al., 1998). Hemogenic potential in the dorsal aorta can be readily identified by expression of endothelial markers (CDH5, CD31, ACE) and nascent transcription factors such as Gata2, Runx1, and Gfi1 (Fadlullah et al., 2021; Gao et al., 2013; Minegishi et al., 1999; Rybtsov et al., 2011; Taoudi and Medvinsky, 2007; Taoudi et al., 2008; Thambyrajah et al., 2016; Tsai et al., 1994). Co-expression of several surface markers and transcription factors are associated with HSC activity, including Sca1, Gpr56, CD27 and EPCR (CD201/PROCR) and multiple signaling pathways have been identified as players of HSPC development (Dzierzak and Bigas, 2018). Next to transcription factors, cytokines and their downstream inflammatory signaling pathways also contribute to HSPC formation, including NF-κB (Espin-Palazon et al., 2014; Li et al., 2014; Sawamiphak et al., 2014). IκBα (NFKBIA) has a well-established role as a repressor of NF-κB factors such as p65 (RELA) and p50 (NFKB1), however IκBα itself can exert gene regulation by direct interaction with chromatin. Chromatin-bound IκBα regulates stem cell specific genes in association with elements of the polycomb repression complex 2 (PRC2), which catalyzes the histone mark H3K27me3, at specific genetic loci (Brena et al., 2020; Marruecos et al., 2020; Mulero et al., 2013). From the AGM, the first HSCs migrate to the FL by E12.5 to amplify and acquire an adult HSCs phenotype (Mahony and Bertrand, 2019). Recent studies indicate that (LT) HSC and HPCs fate is already segregated in the AGM, and that the fetal liver merely serves as a niche for their modest amplification as separate populations (Patel et al., 2022; Yokomizo et al., 2022; Ganuza et al, 2022). Within this context, LT-HSCs are generated as a safeguard pool of cells from the first stages of development, but signaling pathways that allow this segregation and how these LT-HSCs protect their integrity during development remains unclear. In the adult bone marrow, LT-HSC are characterized as LSK CD48-CD150+ with further classification as dormant, the most naïve state of HSCs, or active, ready for cycling. Dormancy is a mechanism that cells use to preserve their integrity, especially during non-favorable conditions. The dormant LT-HSCs divide slower, and have delayed repopulation/activation capacity when transplanted, although they have an extraordinary capacity of regenerating the hematopoietic system under stress conditions. To protect their integrity, they return to a dormant state once homeostasis of the blood system is re-established (Wilson et al, 2008, Pietras et al, 2014).

The main known regulator that acts as a switch between these states is retinoic acid signaling (RA) (Cabezas-Wallscheid et al., 2017; Kaufmann et al., 2021), next to IFNα (Collins et al., 2021; Essers et al., 2009; Trumpp and Essers, 2012). Intriguingly, RA has been linked to HSC activity in the AGM, and also LT-HSC mobilization in the adult bone marrow (Chanda et al., 2013; Goldie et al., 2008; Purton, 2007; Purton et al., 2000), indicating that signals that regulate the pool of inactive HSC in the adult are already established in the AGM.

Here, we uncover a specific reduction of LT-HSCs in the IκBα knockout new-born pups. This defect is manifested at the FL stage already, and traceable to the first emerging HSCs in the E11.5 AGM, without affecting the general HPC population. We present data indicating that IκBα exerts a PRC2-dependent function in this system, which is comparable to that previously demonstrated in the stem cells of skin and intestine (Marruecos et al., 2020; Mulero et al., 2013). In agreement with this, we detected a decrease of H3K27me3 at the promoters of rarα and rarγ, which is associated with an increase in their transcription. Molecularly and functionally, the IκBα KO LT-HSCs display a dormant (less proliferative) phenotype that is consistent with a delayed chimerism in the recipient mice after transplantation, and is rescued (can be mobilized into cycling) by treatment with a rarα specific inhibitor. We propose that nuclear IκBα regulates self-renewal of LT-HSCs during periods of stress conditions. In its absence, RA levels remain elevated and prevent LT-HSCs to be activated.

Overall, we identify nuclear IκBα as an essential regulator of rarα receptor from the onset of HSC generation in the AGM, which is shifted to rarγ at later stages. This regulatory axis controls the balance between HSC proliferation and dormancy.

## Results

### Inflammatory NF-κB signaling is significantly enriched throughout HSC ontogeny

We set out to identify the inflammatory signaling pathways that are active throughout HSC ontogeny. We therefore analyzed a publicly available dataset that consists of single cell sequencing data of HSC at different stages of development, ie AGM, FL and bone marrow. Non-linear dimension reduction test (UMAP) of all cells showed clustering according to their cell fate, with endothelial cell (EC) situated at the opposing end to bone marrow HSCs (Figure 1A). In between these two fates, the T1/T2 HSC of the AGM and the FL HSCs form the continuum towards bone marrow HSCs (Figure 1A). We reviewed these different HSC populations specifically for inflammatory signaling by gene signature correlation (Figure 1B, suppl table T1) and found its enrichment to increase from early HSC stages to bone marrow LT-HSCs (Figure 1B). Interestingly, the inflammatory pathway NF-κB signaling activity and NF-κB signaling members themselves were increasingly enhanced from T1/T2 HSC to bone marrow HSC fate (Figure 1B and suppl table T1). We further verified the accumulation of NF-κB signaling activity by performing GSEA enrichment analysis either with T1/T2 HSCs or FL-HSCs (Figure 1C). In both instances, the hallmark “Tnfa_signalling_via_ NF-κB” was enhanced in the HSC population compared to EC, with reaching statistically significance in T1/T2 HSCs (Figure 1C). Next, we explored the expression profile of the main NF-kB signaling members across the UMAP (Figure 1D). We found several of them expressed in the different HSC population, including Nfkbia (hereafter IκBα) (Figure 1D and suppl Figure S1). IκBα is the main regulatory factor of NF-κB signaling and additionally, it can also regulate stem cell specific genes in association with the PRC2 complex (Mulero et al., 2013). We therefore assessed the impact of global IκBα loss during hematopoietic development.

**Figure 1:**
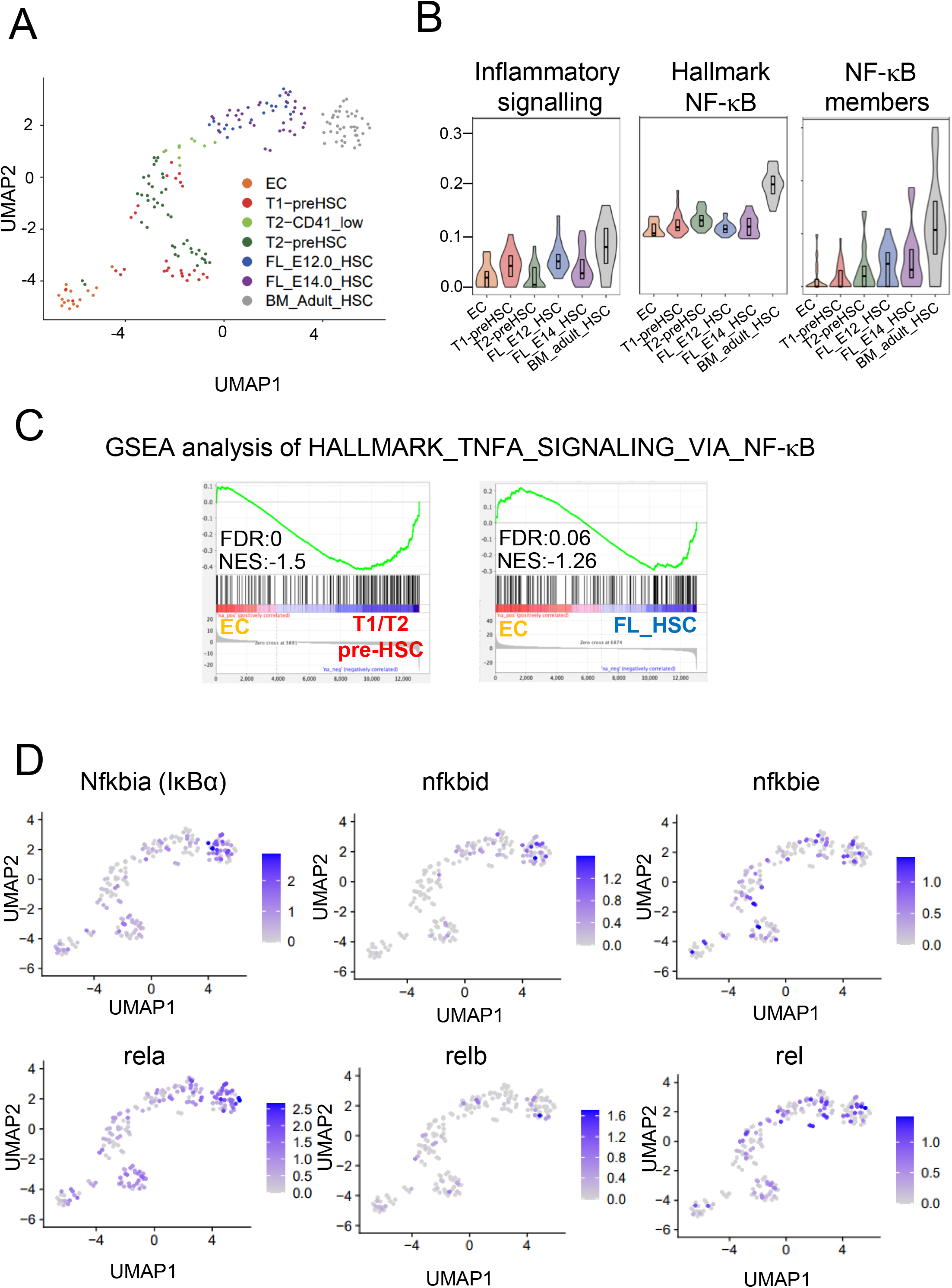
NF-κB signaling is enriched in AGM derived HSCs. (**A**) UMAP based clustering of the single cells based on gene expression with colors highlighting the different HSC populations. Single cell RNA sequencing data derived from Zhou et al, 2016. (**B**) enrichment score analysis of indicated HSC populations for inflammatory, NF-κB pathway and NF-κB signaling molecules. (**C**) GSEA plots comparing endothelial cells (EC) with either T1/T2 HSC or FL-HSC for genes in the Hallmark “TNFA_SIGNALING_VIA_ NF-κB”. (**D**) UMAP representation of NF-κB signaling molecule gene expression in the indicated HSC population.

### IκBα loss compromises the generation of cKIT+CD45+Gfi1+EPCR+ HSC specific subpopulations in the AGM

First, we performed Immunohistochemistry (IHC) on E11.5 AGM sagittal sections of *Gfi1:tomato* embryos for total IκBα (Figure 2A, Suppl Fig S2A). We discovered accumulation of a punctuated nuclear signal for IκBα mainly in *Gfi1* positive HE and IAHC (Dll4 is a marker for DA, Figure 2A, Suppl Fig S2A). We further validated our findings with a second strategy by staining ckit positive IAHC cells with an antibody that recognizes the phosphorylated form of IκBα (p-IκBα Ser 32,36) that has been linked to nuclear localization and stem cell function (Mulero et al., 2013; Marruecos et al., 2020). We detected nuclear p-IκBα staining mainly in IAHC (Fig 2B) with heterogeneous distribution in around 50% cKIT positive cells (Fig 2C). Next, we assessed if the presence of IκBα in a subset of IAHC had a functional impact. We therefore examined the hematopoietic sub-populations within the E11.5 AGMs of WT, or with global loss of IκBα by FACS analysis. The frequency of the overall CD31+cKIT+/CD45+ (HSPCs) was not altered in IκBα KO embryos when compared with the controls (Figure 2Di, Suppl Fig S2B). However, inclusion of additional markers that restrict this population further to HSCs by sub-gating for GFI1 and EPCR (Dignum et al., 2021; Hadland et al., 2022; Zhou et al., 2016b) led to the detection of a significant decline in the frequency of HSCs in the IκBα KO (Figure 2Dii, Suppl Fig S2B). Importantly, NF-κB activity was not substantially increased in the IAHC/HSC after IκBα removal as indicated the primary cytoplasmic p65-NF-κB localization in the AGM region of IκBα KO (comparable to the WT) (Suppl Fig S2C).

**Figure 2:**
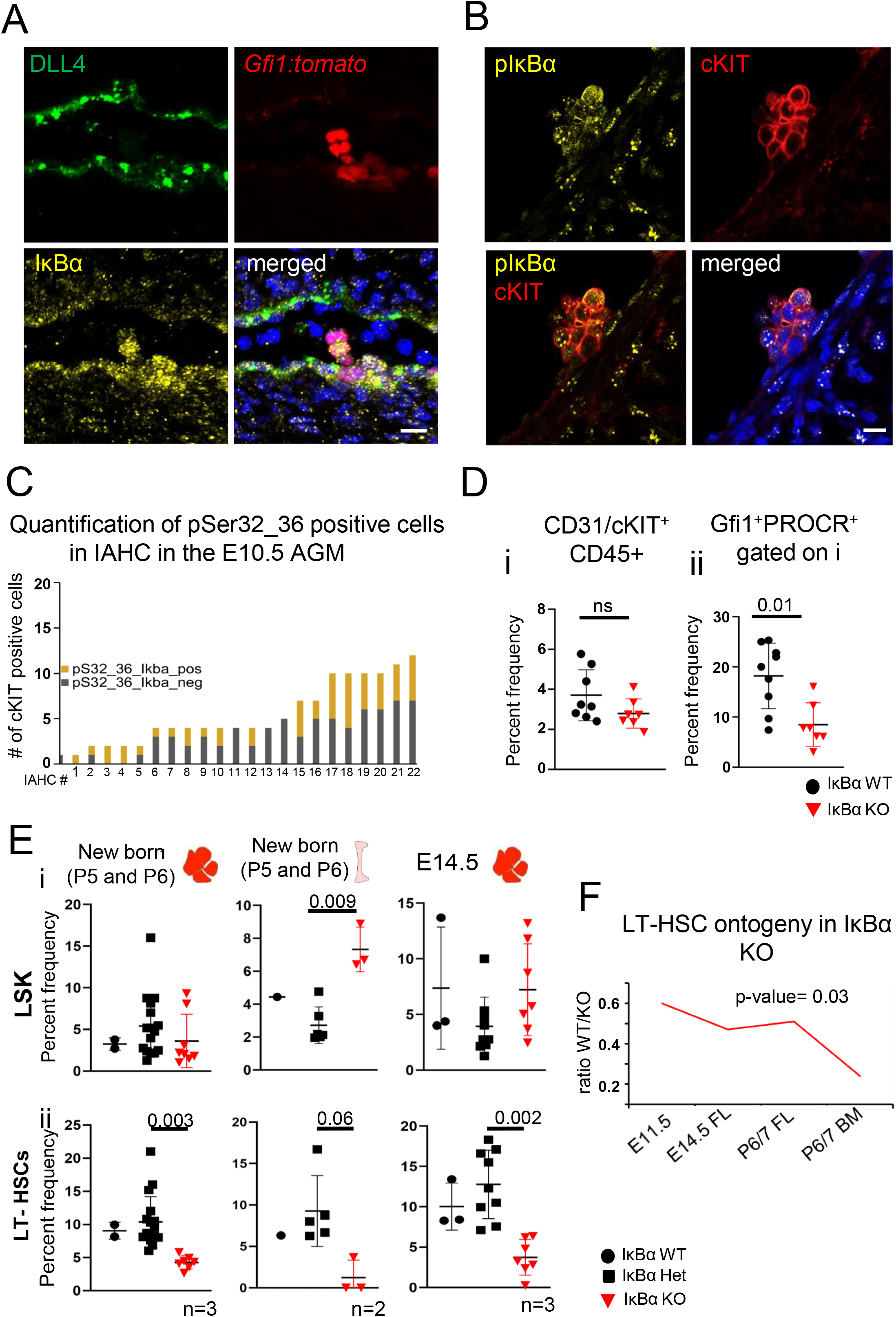
Nuclear IκBα is expressed in Gfi1^+^ HE/IAHC and its loss in the AGM reduces (LT-) HSCs. (**A**) IHC on E11.5 (43-45s) sagittal AGM section for DLL4 (green) and I IκBα (yellow) and DAPI (blue) on *Gfi1:tomato* (red) transgenic embryos. Scale bar: 20μm. (**B**) IHC on E11.5 (42-45s) cross section of AGM for pSer32_36 IκBα (yellow), cKIT (red) and DAPI (blue). Scale bar: 10μm. (**C**) Bar chart depicting the number of pIκBαSer32_36 positive cells within cKIT+ IAHC. Each bar represents one cluster. (**D**) Bar chart depicting individual values for the frequency of (**i**) cKIT^+^CD45^+^(IAHC) in CD31+ cells and (**ii**) Gfi1^+^EPCR^+^ positive cells within IAHC of E11.5 (42-46s) AGMs of IkBα WT and KO, n=15 AGMs in 3 independent experiments. P-value determined with two tailed t-test. (**E**) Bar chart with individual values for the frequency of (**i**) LSK (lin-Sca+cKIT+) liver of newborn (P5/6) IkBα WT, het and KO, n=26 pups in 3 independent experiments, frequency of LSK in the bone marrow of newborn (P5/6) IkBα WT, het and KO, n=9 pups in 2 independent experiments and values for the frequency of LSK in the foetal liver of E14.5 IkBα WT, het and KO, n=19 pups in 3 independent experiments. (**ii**) Bar chart with individual values for the frequency of LT-HSCs (LSKCD48-CD150+) in the liver of newborn (P5/6) IkBα WT, Het and KO, n=26 pups in 3 independent experiments, and individual values for the frequency of LT-HSCs in the bone marrow of newborn (P5/6) IkBα WT, het and KO, n=9 pups in 2 independent experiments in the foetal liver of E14.5 IkBα WT, het and KO, n=19 pups in 3 independent experiments. P-values determined with two tailed t-test with graphpad prism8. (**F**) Chart with the calculated ratios of the average (LT-) HSC frequency from the E11.5 AGM, E14.5 FL and P6/7 newborn foetal liver and bone marrow (taken from Fig 2Eii) in IkBα KO over WT. P-values determined with two tailed t-test with graphpad prism8.

### Reduced number of LT-HSC but normal lineage differentiation in IκBα KO fetal liver and new-born pups

To examine if this decline in the AGM HSC population had a knock-on effect later in development, we analyzed the frequency of the HSCs in the FL at E14.5 (where HSCs expand and mature) and in newborn pups (Suppl Figure S3A), since IκBα deficiency leads to death beyond 7 days after birth (Beg and Baldwin, 1993). All blood lineages were present in the IκBα KO new-borns, although we detected a significant reduction in the B220 cell compartment in the bone marrow and in the liver (Suppl Fig S3B), as previously reported (Feng et al., 2004; Liou and Hsia, 2003). The frequency of the LSK was comparable between the genotypes in the E14.5 fetal liver, bone marrow and new-born liver (Figure 2Ei, Suppl Fig S3C), but the number of LT-HSCs was significantly reduced, in all hematopoietic sites of IκBα KO and the difference in the number of LT-HSCs between WT and KO increased during hematopoietic development (Figure 2Eii, 2F and Suppl Fig S3C) with no significance between IκBα WT and HET.

### FL LT-HSCs retain an AGM specific gene expression profile in the absence of IκBa

To capture the molecular alterations precisely in the E14.5 FL LT-HSCs of IκBα KO, we sorted 500 WT/HET or IκBα KO LSK/CD150+CD48-by FACS and performed bulk RNA seq analysis of the samples (Fig 3A). Principal component analysis (PCA) showed a separation of the samples depending on their genotype with the maximum differences observed in the KO cells (Suppl Fig S4A). Specifically, we found 204 and 365 significantly up and down regulated genes respectively, between IκBα WT/HET and KO (Fig 3B, Suppl Table T2). We first investigated whether IκBα loss increased NF-κB signaling in this compartment by comparing gene expression level of known NF-κB target genes (Suppl table T2, (Pahl, 1999) including IRF1/2, A20 (TNFAI), Dctn3 and Sqstm1. Although specific genes such as IRF1 and 2 were upregulated in the IκBα KO LT-HSCs, overall, their expression levels were not consistently elevated between the genotypes (Suppl Fig S4B, Suppl table T2), further indicating that the NF-κB pathway was not hyper activated in the absence of IκBα. A functional enrichment analysis using KEGG database showed that ribosomal biogenesis, phosphatidylinositol (PI) signaling system, and Insulin resistant related genes were deregulated in the FL LT-HSC of IκBα KO (Suppl Fig S4C; Suppl table T2). Crucially, we detected higher expression of critical embryonic AGM HSC associated signaling pathways, including Interferon alpha, IL6/JAK/STAT, Wnt and TGFβ signaling in IκBα KO (Suppl Fig S4D). We therefore examined the expression levels of key embryonic (AGM) HSC-associated genes such as Smad6, Sox18, Gpr56, gata2 and Neurl3 in our data set. We found higher expression levels of all tested AGM HSC associated genes in the IκBα KO FL cells (Fig 3C). These results suggested that IκBα deficiency imposes a defect in the development of HSCs from the AGM to the FL stage. To test this hypothesis, we performed GSEA of genes upregulated in IκBα KO LT HSC against two data sets; one dataset consists of the top 100 genes up- or down regulated from hemogenic endothelium and HSCs of our (unpublished) single cell RNA seq (Fig 1C) and are therefore characteristic for nascent AGM HSCs. The second dataset comprises a FL LT-HSC signature (Manesia et al., 2017). Strikingly, the IκBα KO LT-HSC correlate at a much higher degree, and significantly with the AGM-associated HSC profile (NES= - 2.63), than with the FL LT-HSC signature (NES= -1.16) (Fig 3D, Suppl Fig S4E), indicating that IκBα KO FL LT-HSCs are retained in a previous developmental AGM stage.

**Figure 3:**
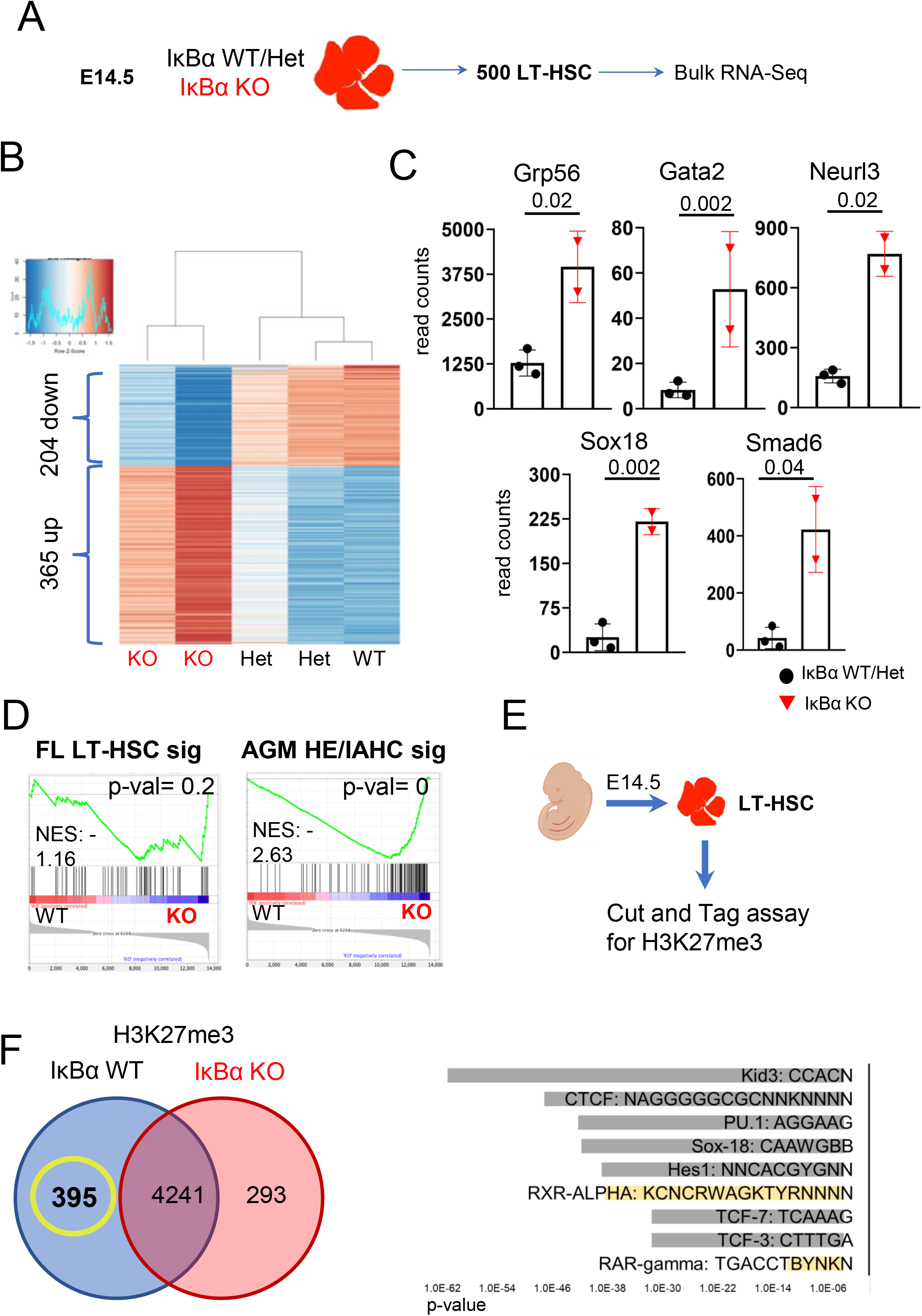
IκBα KO LT-HSCs maintain an AGM specific HSC gene expression program. (**A**) Scheme of 500 LT-HSC purification for bulk RNA seq from IκBα WT, Het or KO embryos at E14.5. replicates from 2 independent experiments. (**B**) Heat map of differentially expressed genes (</>2log2) between IκBα WT/Het and KO. (**C**) Scatter plot with the reads counts in IκBα WT/Het and KO LT-HSCs for selected genes that are commonly linked to AGM hematopoiesis HSCs. P-values determined with two tailed t-test with graphpad prism8. (**D**) GSEA plots from comparisons of genes upregulated in IκBα KO in LT-HSC from the RNA seq and signatures for fetal liver LT-HSCs (left) and AGM HSCs (right). (**E**) Schematic representation of the workflow for cut and tag assay on sorted LT-HSC from E14.5 FL. (**F**) Venn diagram with number of peaks/promoter regions obtained from cut and tag for H3K27me3 in LT-HSC and comparing IkBα WT and KO. Peaks derived from 3 independent experiments and at least present in 2 replicates. (G) Bar chart showing the transcription factor motifs and their p values associated with H3K27me3 peaks that are unique to IκBα WT.

### Retinoic acid signaling is increasingly activated in the AGM and FL LT-HSCs of IκBα-deficient embryos

Next, we aimed to investigate whether IκBα was specifically modulating genomic loci in LT-HSCs in a PRC2 dependent manner as previously described (Mulero et al., 2013). In order to identify the target genes that are subject to the IκBα-dependent deposition of H3K27me3, we used cut and tag assay on sorted LT-HSC from WT and KO E14.5 FL (Fig 3E) and compared the differentially enriched peaks. Intriguingly, 293 and 395 peaks were uniquely detected in the IκBα KO or WT LT-HSC, respectively. Whilst the peaks in the KO suggest less accessible loci in WT (Suppl Fig S5A), unique peaks in the WT indicate higher chromatin accessibility and altered gene expression from these loci in IκBα KO LT-HSCs. These genes were significantly enriched in calcium signaling and signaling by GPCR (Suppl table T3). We further explored if any transcription factor binding motifs were overrepresented within these hypo methylated (more accessible) regions in the IκBα KO LT-HSCs. Next to CTCF, Pu1, TCF3 and TCF7, we discovered a significant enrichment of the retinoic acid receptor binding motif (Fig 3F and Suppl table T3), suggesting that retinoic acid signaling may be over activated in the IκBα KO LT-HSCs.

### Increased activation of retinoic acid receptor rarα in the AGM, and rarγ in the FL LT-HSCs of IκBα-deficient impose a dormancy signature

Next, we performed cut and tag assay with anti-IκBα and H3K27me3 antibodies on sorted LT-HSC and isolated CD31 positive AGM cells (Fig 4A). Although cut and tag assays for low cell numbers are challenging, we found a reproducible peak for IκBα in the promoter of rarγ and not rarα in LT-HSC that coincides with a region containing reduced levels of H3K27me3 in the IκBα KO LT-HSC (Fig 4B, Suppl Fig S5B). In the AGM-derived CD31+ cell cut and tag assay however, IκBα itself showed a peak at the promoter region of only rarα, the pivotal receptor for HSC function in the AGM (Fig 4C, Suppl Fig S5B, Suppl table T3) (Chanda et al., 2013). We then examined the levels of rarα and rarγ expression in our LT-HSC RNA seq data set. Consistent with our cut and tag data, we found significantly increased levels of rarα in IκBα KO LT-HSC (Fig 4D) and a trend for elevated levels of rarγ (data not shown).

**Figure 4:**
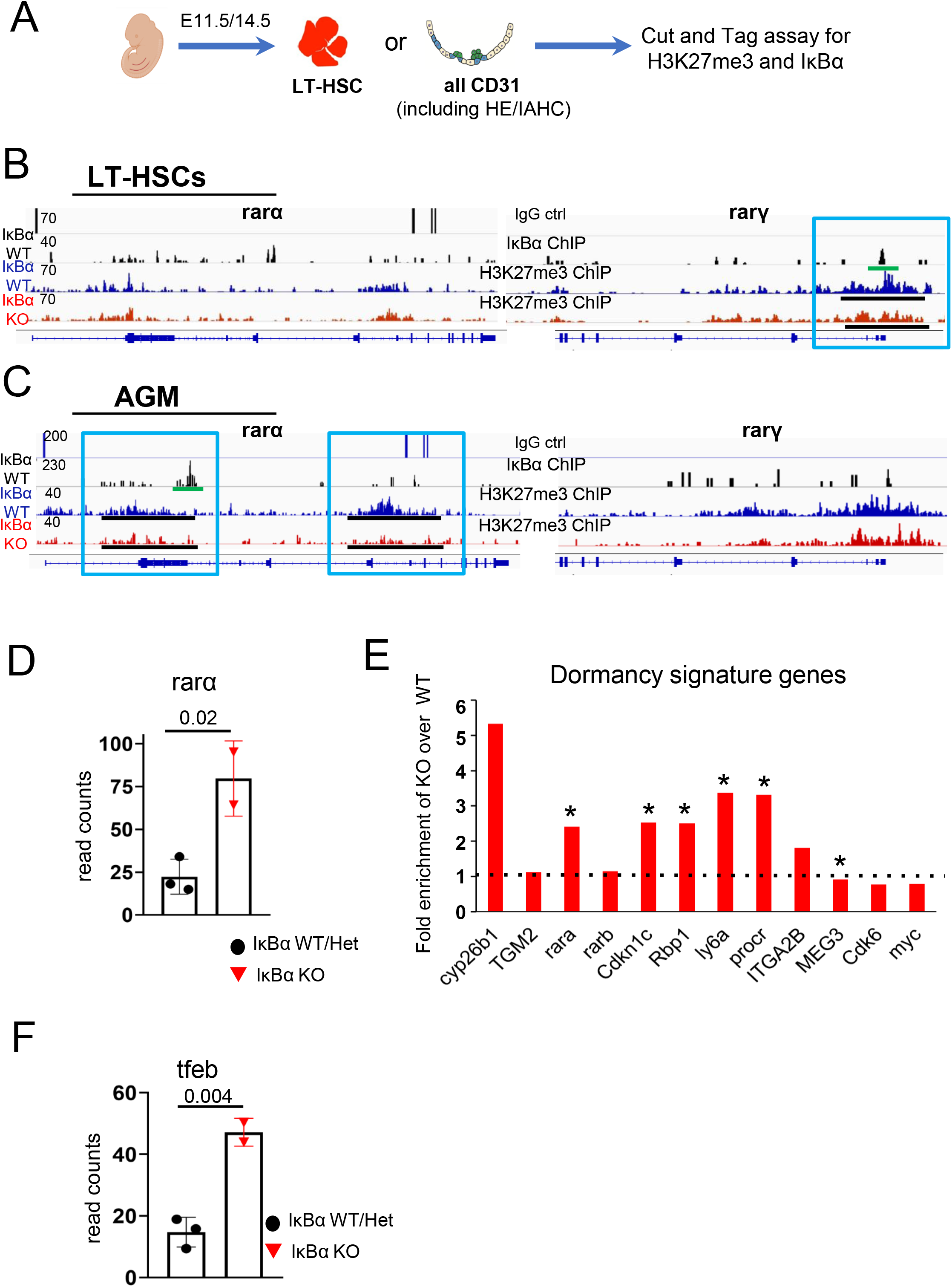
IκBα binds to the retinoic acid receptors rxrα, rarα and rarγ and induce dormancy. (**A**) Schematic representation of the workflow for cut and tag assay on sorted LT-HSC from E14.5 FL and sorted CD31+ cells from E11.5 AGMs. (**B**) Snapshots from igv browser depicting enrichments obtained from cut and tag assays with LT-HSC of E14.5 foetal liver at the promoters of the rarα and rarγ genes with H3K27me3 (relevant peak highlighted with a black bar) and IκBα antibodies (relevant peak highlighted with green bar). 2 independent replicates for H3K27me3 and 1 for IκBα. (**C**) Snapshots from igv browser depicting enrichments obtained from cut and tag assays with CD31+ AGM cells at the promoters of the rarα and rarγ genes with H3K27me3 (relevant peak highlighted with a black bar) and IκBα antibodies (relevant peak highlighted with green bar). One replicate for H3K27me3 and 2 independent replicates for IκBα. (**D**) Scatter plot with the reads counts for rarα from IκBα WT/Het and KO LT-HSCs RNA seq data set. (**E**) Bar chart showing the read count differences in IκBα WT/Het and KO for dormancy related genes. (**F**) Bar chart showing the read count differences in IκBα WT/Het and KO for tfeb. Reads values taken from the RNA seq of E14.5 LT-HSCs. P-values determined with two tailed t-test with graphpad prism8.

Since increased retinoic acid signaling is associated with dormancy of LT-HSCs (Cabezas-Wallscheid et al., 2017; Schönberger et al., 2022), we cross-referenced the published list of key dormancy-related genes in LT-HSC with our RNA-seq data set (for genes differentially expressed in IκBα genotypes). We found several of these key genes such as p57, Tgm2, Rbp1, Cyp26b1 and Ly6a up-regulated and myc and cdk6 down regulated in IκBα KO, a pattern compatible with LT-HSC in a dormant state (Fig 4E). Further supporting the molecular features of dormant cells, the IκBα KO LT-HSCs also express significantly higher levels of the transcription factor EB (tfeb) (Figure 4F), whose elevated levels is a further indicator of dormant cell (García-Prat et al., 2021; Garcia Prat et al., 2020).

Altogether, our data indicates an over activation of the retinoic acid signaling in the IκBα-deficient LT-HSC that is initiated already in the AGM, and the high retinoic acid signaling levels are maintained in the IκBα-deficient FL LT-HSC and in its most naïve sub-population leading to an increased level of dormancy at the molecular level.

### IκBα-deficient LT-HSCs are functionally dormant

Finally, we tested IκBα-deficient LT-HSCs for their retinoic acid-dependent dormant phenotype. In the first instance, we determined their proliferation potential *in vivo*. Pregnant females with E13.5 embryos were intra peritoneally injected with 2mg of BrdU (Fig 5A). After 2 hours, the females were sacrificed to analyze the cell cycle status of FL LT-HSCs from WT and KO embryos by flow cytometry. We observed a significantly higher proportion of LT-HSC that had not incorporated BrdU in the IκBα -deficient embryos (Fig 5A), a clear indication that these LT-HSC were cycling less. We then explored if the inhibition of the retinoic acid signaling was sufficient to rescue the non-cycling status of the IκBα -deficient LT-HSC. We therefore purified lineage-depleted E14.5 FL cells from IκBα WT or KO embryos and treated these cells with either control or 1μM of the rarα specific inhibitor Ro-415253 (hereafter Ro-41) for 2 days. The cells were then examined for their cell cycle profile (Fig 5B). We found that treatment with Ro-41 increased the frequency of cycling cells both in the WT and IκBα KO LT-HSCs. Notably, treating IκBα KO LT-HSCs with a rarα specific inhibitor was sufficient to reduce the number of dormant (G0) cells to the levels of the IκBα WT (Fig 5B), supporting the concept that increased retinoic acid signaling is a causal mediator of cellular dormancy in the IκBα LT-HSCs. Dormant HSC shows a paradox behavior upon stress stimuli, including transplantation assays or *ex vivo* culture. In these conditions, the cells become activated with a delay, but outperform their WT counterpart over time (Cabezas-Wallscheid et al., 2017; Kaufmann et al., 2021) since these cells need more time to become activated and go into cycle, but at the same time possess a higher self-renewal and repopulation capacity than their activated HSC counterparts (Viale and Pelicci, 2009). We therefore tested the hematopoietic activity of IκBα KO LSK (1000 cells) in 1:1 competition with WT LSK cells in transplantation assay (Fig 5C). Indeed, PB blood analysis 4 weeks after the transplantation showed a significant lower blood chimerism from IκBα KO LSK cells. But in the following bleedings (8, 12 and 16 weeks), the IκBα KO cells progressively reached a similar level of engraftment to their WT counterparts (Fig 5D). By 4 months after transplantation, there was no significant difference between the genotypes in their contribution to lineage differentiated cells (Fig 5D). All blood lineages were reconstituted by both IκBα WT and KO LSK cells, albeit a small increased contribution of the IκBα KO LSK cells to the B cell lineage (Suppl Fig S6B). Additionally, we also assayed E11.5 IκBα WT or KO AGMs for stem cell activity. Here, 1.75ee were transplanted per recipient and after 4 months, 4/4 WT and 2/3 IκBα KO recipients showed donor chimerism >5% in the bone marrow (Suppl Fig S6C). In both assays we conducted secondary transplantations (Fig 5E and Suppl Fig S6D) and in both instances, we detected a trend to higher repopulating capacity of IκBα KO cells when considering > 5% bone marrow chimerism (Fig 5E), indicating that in the absence of IκBα, activated LT-HSCs cannot return to a dormant state.

**Figure 5:**
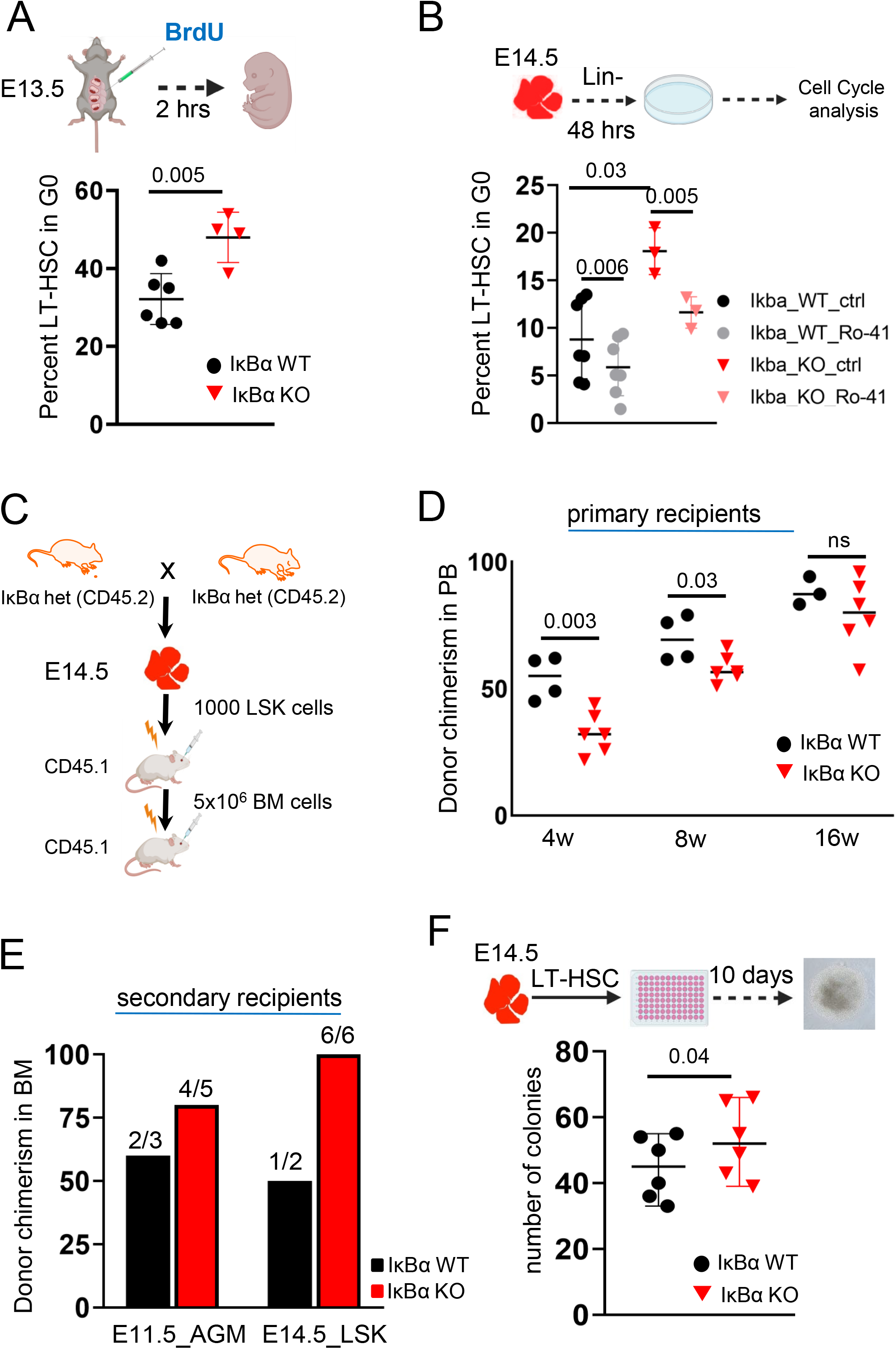
IκBα KO (LT)-HSC are functionally dormant and can be mobilized by rarα inhibition. (**A**) Bar chart with the frequencies of LT-HSC in G0 phase of cell cycle after 2 hrs of BrdU labeling *in vivo* in E13.5 IκBα WT or KO FL. Data originates from 2 independent experiments with 10 embryos. P-value determined with two tailed t-test. (**B**) Bar chart with the frequencies of LT-HSC in G0 phase of cell cycle after 48hrs of *ex vivo* culture of E14.5 IκBα WT or KO FL cells treated with 10uM of Ro-41 or DMSO. Data originates from 3 independent experiments with 10 embryos. P-value determined with two tailed t-test. (**C**) Scheme of transplantation experiment set up. (**D**) Bar chart with the frequencies of donor chimerism (CD45.2) in the peripheral blood of primary recipient (CD45.1) in 4-week intervals. Data originates from one experiment with 10 embryos. P-value determined with two tailed t-test. (**E**) Bar chart with the frequencies of donor chimerism in the bone marrow of secondary recipient at 16 weeks. Data originates from two experiments with 18 mice. P-value determined with two tailed t-test. (**E**) Bar chart with the number of hematopoietic colonies from LT-HSC deposited as single cell after 10 days in culture from IκBα WT or KO E14.5 FL. Data originates from 3 independent experiments with 6 embryos. P-value determined with two tailed t-test.

We further assessed the *in vitro* clonogenic activity of IκBα WT and KO LT-HSC specifically. In support of enhanced LT-HSC activity of IκBα KO cells over time and in stress conditions, we found more of the IκBα KO LT-HSC forming colonies when they were sorted as single cells into wells (Fig 5F). An increased proliferation rate upon stress was also detected by CFSE labelled cell proliferation assay (data not shown).

Altogether, we show that IκBα deficiency imposes an increased dormant phenotype in the LT-HSC population under physiological conditions. Upon stress the IκBα LT-HSCs become activated but cannot return to a dormant state, consistent with previous reports (Cabeza-Wallscheid et al, 2017).

## Discussion

NF-κB is a pleiotropic transcription factor that responds to inflammation by activating multiple biological processes in emergency situations. Inflammatory signals are in turn coordinated with cellular differentiation, however the mechanisms underlying this crosstalk are not well understood. In the embryo, the environment is generally protected from insults. Nevertheless, inflammatory signals, including canonical NF-κB signaling are critical for HSPC formation in the AGM (Espin-Palazon et al., 2014; Mariani et al., 2019), inferring that there must be safe guards to protect LT-HSCs from extensive differentiation and self-renewal. Our data show that nuclear IκBα activity could be a deterministic factor for LT-HSCs response to inflammation already in the AGM. Here, it primes the small pool of LT-HSCs for self-renewal and proliferation via reducing the levels of RA signaling in these cells. Of note, recent studies have established that all HSC activity is contained within RA metabolizing cells (Chanda et al, 2013; Schonberger et al, 2022; Luff et al, 2022). Although the main function of IκBα is the inactivation of NFκB complex by cytoplasmic retention or by active removal from the nucleus (Beg and Baldwin Jr., 1993), there is an ancestral function of IκBα that is evolutionary conserved and is directly affecting PRC2 complex activity on stem cell specific genes (Brena et al., 2020; Mulero et al., 2013).

Here, we report that IκBα-deficient embryos can specify all blood lineages and progenitors, but they show a significant decrease specifically in the number of HSC generated at different developmental stages, likely due to their premature dormant phenotype which is evident at the molecular level in their expression of dormancy signature genes. Moreover, this HSC pool is already compromised at the site of their origin in the E11.5 AGM since we find that the low number of IκBα KO LT-HSC that are present in the FL at E14.5 retain embryonic/AGM specific signature, including Sox18, Gpr56 or GATA2, indicating that the dormant phenotype prevented them from self-renewal in the FL. Functionally, these LT-HSCs cycle less *in vivo*, but respond with delayed activation to stress. These observations agree with previous findings in the IκBα-deficient adult intestinal stem cells that maintain a fetal signature (Marruecos et al., 2020). In both scenarios, adult stem cell programs are compromised in the absence of IκBα.

Our results further show that NFκB is not hyper activated in the AGM or FL HSCs in the absence of IκBα and supports the importance of the chromatin function of IκBα. Indeed, nuclear IκBα might modulate further pathways that are important for HSC generation. However, the specific involvement of chromatin IκBα in the LT-HSC specification is still challenging to assess since we need to determine IκBα (or other factors) binding across the genome in small populations. We have therefore probably not detected many of the IκBα-bound genes. Using cut and tag experiments for H3K27me3 and IκBα itself, we reveal direct binding of IκBα to the promoter of the rarα in the HSPC-containing CD31+ cells of the AGM. Indeed, RARα induction through stimulation with the rarα specific agonist AM580, or its inactivation through RO-415253 has been irrefutably linked to HSC emergence and activity in the AGM (Chanda et al., 2013; Goldie et al., 2008; Marcelo et al., 2013).

Strikingly, we observe a switch in the retinoic acid receptor isoform expression between the AGM and FL. In the AGM derived cut and tag for H3K27me3, we detect less of this repressive mark in the promoter of rarα in IκBα KO, along with accumulation of IκBα itself in WT cell, indicating that this receptor is under direct regulation by IκBα while the rarγ promoter region is unchanged for H3K27me3 and has no peak for IκBα. This scenario is reversed in the FL LT-HSCs where we detected less H3K27me3 in IκBα KO HSCs and a specific peak for IκBα in the rarγ promoter of WT FL LT-HSCs. From studies in the LSK compartment of adult bone marrow, RARγ has been proposed as a regulator of stem cell renewal (Purton, 2007; Purton et al., 2000). Recent studies on purified adult LT-HSCs propose that the retinoic acid receptor RARβ is responsible for the switch between dormancy and activation (Schönberger et al., 2022). In our RNA sequencing data from FL LT-HSC, rarβ levels are hardly detected. Based on all this data, we hypothesize that different isoforms prevail at different stages of hematopoiesis. In our scenario, rarα, the HSC regulating isoform in the AGM (Chanda et al., 2013), is maintained at high levels in FL LT-HSCs that would normally switch to rarβ. Under normal physiological conditions, inflammatory signals in the microenvironment would increase the levels of chromatin IκBα and PRC2 activity on the rarα promoter leading to lower RARα expression, and likely signaling to induce self-renewal or differentiation. In the absence of IκBα, the HSCs maintain elevated levels of rarα and present a dormant phenotype. Finally, we cannot exclude that classical NF-κB transcriptional activity is also involved in regulating the HSC quiescence or activation. It is noteworthy that NF-κB regulators may likely control chromatin IκBα, since inflammatory signals such as TNFα regulate chromatin bound IκBα (Mulero et al., 2013). The IκBα pups die from day 7-10 after birth from a p65-dependent severe skin inflammatory reaction (Rebholz et al., 2007). Considering our results, we believe that in the embryo, the inflammatory response due to NF-κB over activation is not initiated; however in conditions where cells are exposed to external stimuli, NF-κB response becomes decisive for survival. Finally, we also detect higher levels of the transcription factor EB (tfeb) in the IκBα KO LT-HSCs. This transcription factor controls the endolysosomal function of dormant HSCs, and its higher levels of expression has been experimentally validated as a hallmark of dormancy (García-Prat et al., 2021; Garcia Prat et al., 2020).

Altogether, we postulate that (LT)-HSCs must interpret the inflammatory signals that they receive in the AGM differently to HPC that leads to the nuclear function of IκBα. But how this difference in nuclear IκBα activity is achieved remain to be understood.

We are yet to fully uncover the incorporation of the several inflammatory signals into a cellular response although our data agrees with the model depicted in Fig 6. It will be intriguing to define the inflammasome that allows the entry and exit out of a dormant state from HSCs which would significantly improve our basis for regenerative medical approaches.

**Figure 6:**
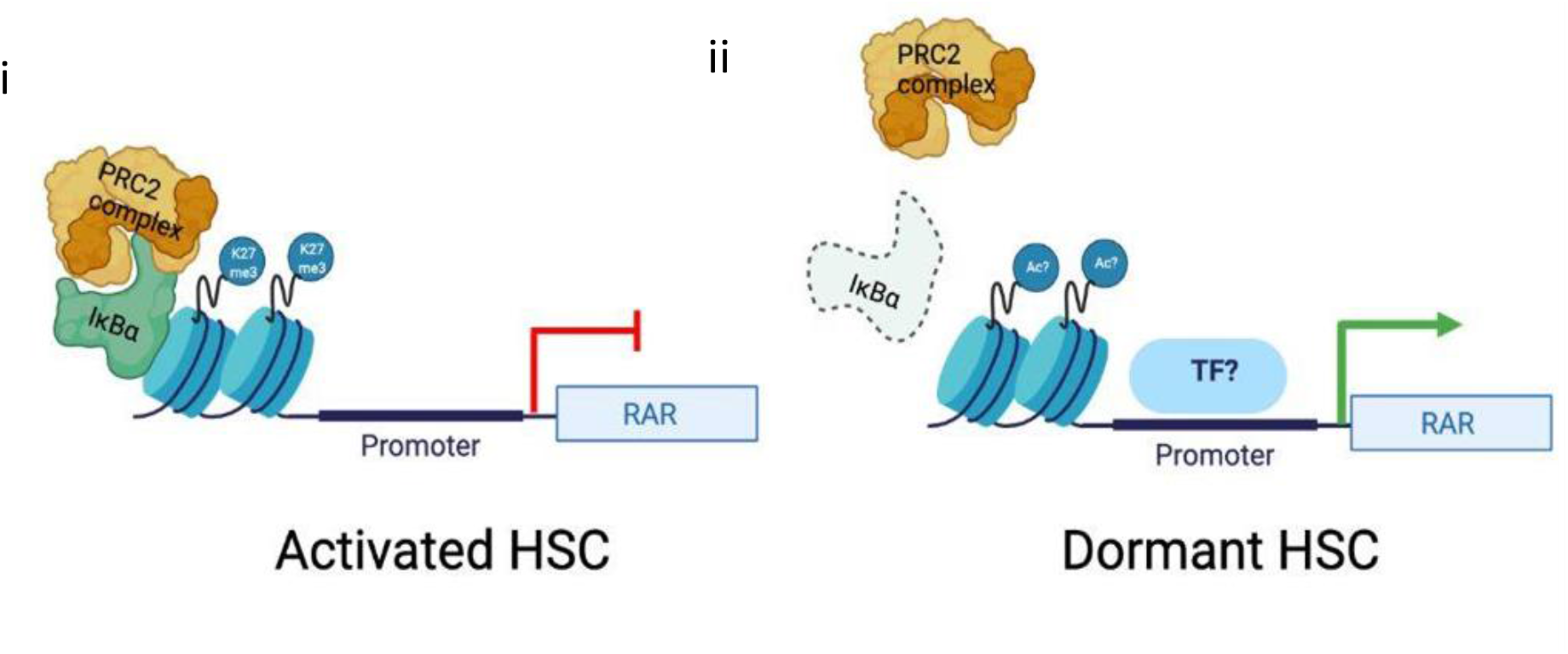
Schematic summary of the main findings. (**i**) IκBα associates with PRC2 complex at RAR gene promoter and silences retinoic acid signaling in the absence of stimuli (**ii**) in the absence of IκBα this silencing is lost and retinoic acid signaling is hyper-activated.

## Data availability

RNA-Seq and cut and tag-seq data: GEO accession number GSE188525.

## Acknowledgements

We thank all members of Espinosa and Bigas laboratories for helpful discussions and technical support. We also thank the animal facility, FACS facility, and genomic facility of the PRBB, CRUK Manchester Institute and the single cell Unit of the IJC for their technical support. This work was funded by grants from SAF2016-75613-R, PID2019-104695RB-I00 and PDC2021-120817-I00 from Agencia Estatal de Investigación (AEI) and SLT002/16/00299 from Department of Health, Generalitat de Catalunya. The work in G.L. laboratory is supported by Blood Cancer UK (19014) and Cancer Research UK Manchester Institute Core Grant (C5759/A27412) RT is a recipient of BP2016(00021) and BP/MSCA 2018(00034) fellowship programs from Generalitat de Catalunya/MSCA.

## Author contributions

AB, RT, and LE conceptualized the study, designed the experiments, analyzed data and wrote the manuscript. RT, WHN, PH, CB, AP, JG, NC, LM and CR-H performed experiments and analyzed data. MP analyzed the cut and tag data. MC_P analyzed the LSK scRNAseq under supervision of EM. YG and ZF analyzed the E11.5 AGM scRNAseq from Zhou et al, 2016 and bulk RNA-sequencing data of LT-HSCs. AB, RT, GL and LE wrote the manuscript.

## Figure legends

**Suppl Figure S1:**
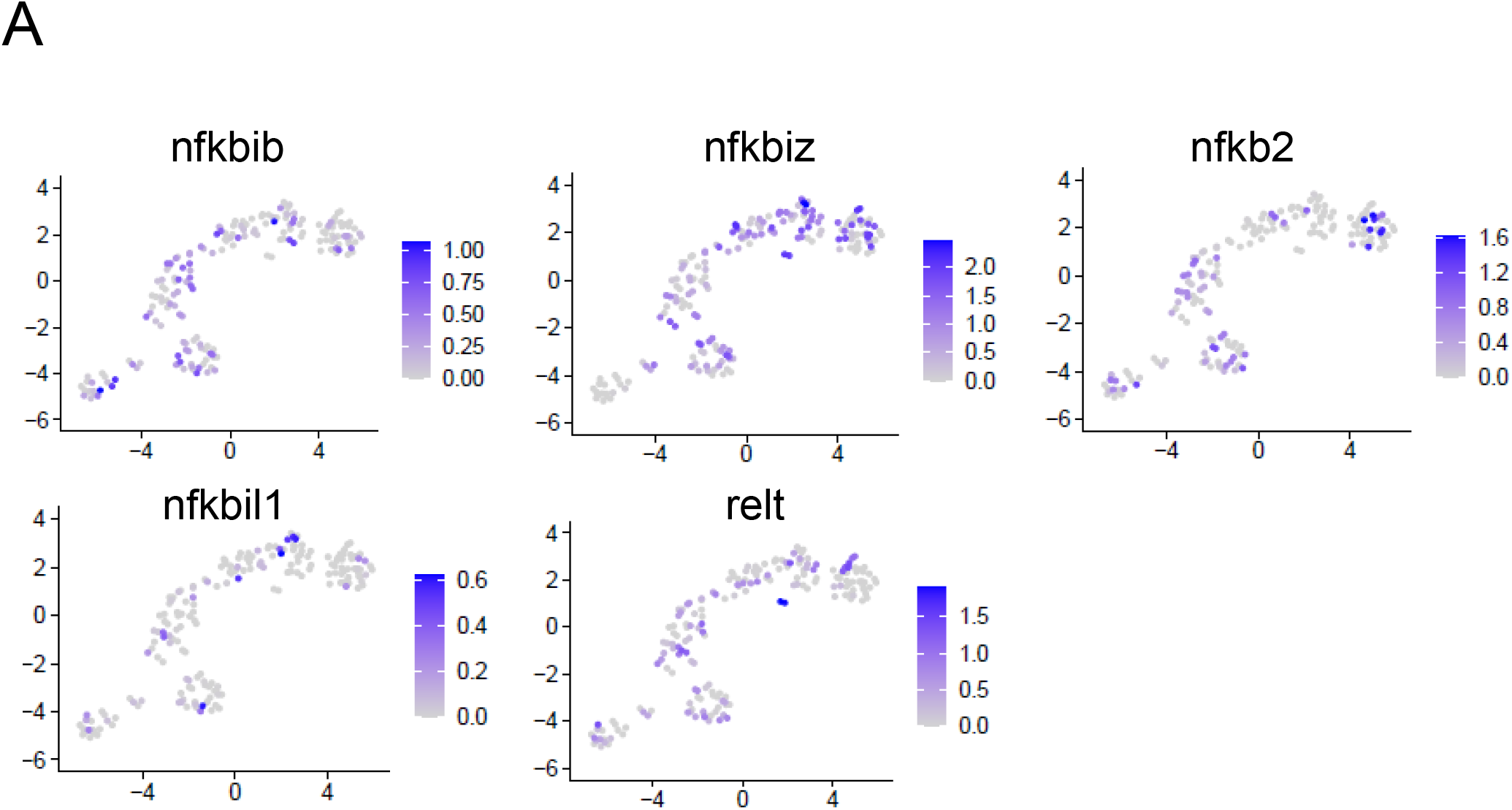
Gene profiling denotes NF-κB signaling presence in HSCs. (**A)** Representation of NF-κB signaling molecules across the UMAP.

**Suppl Figure S2:**
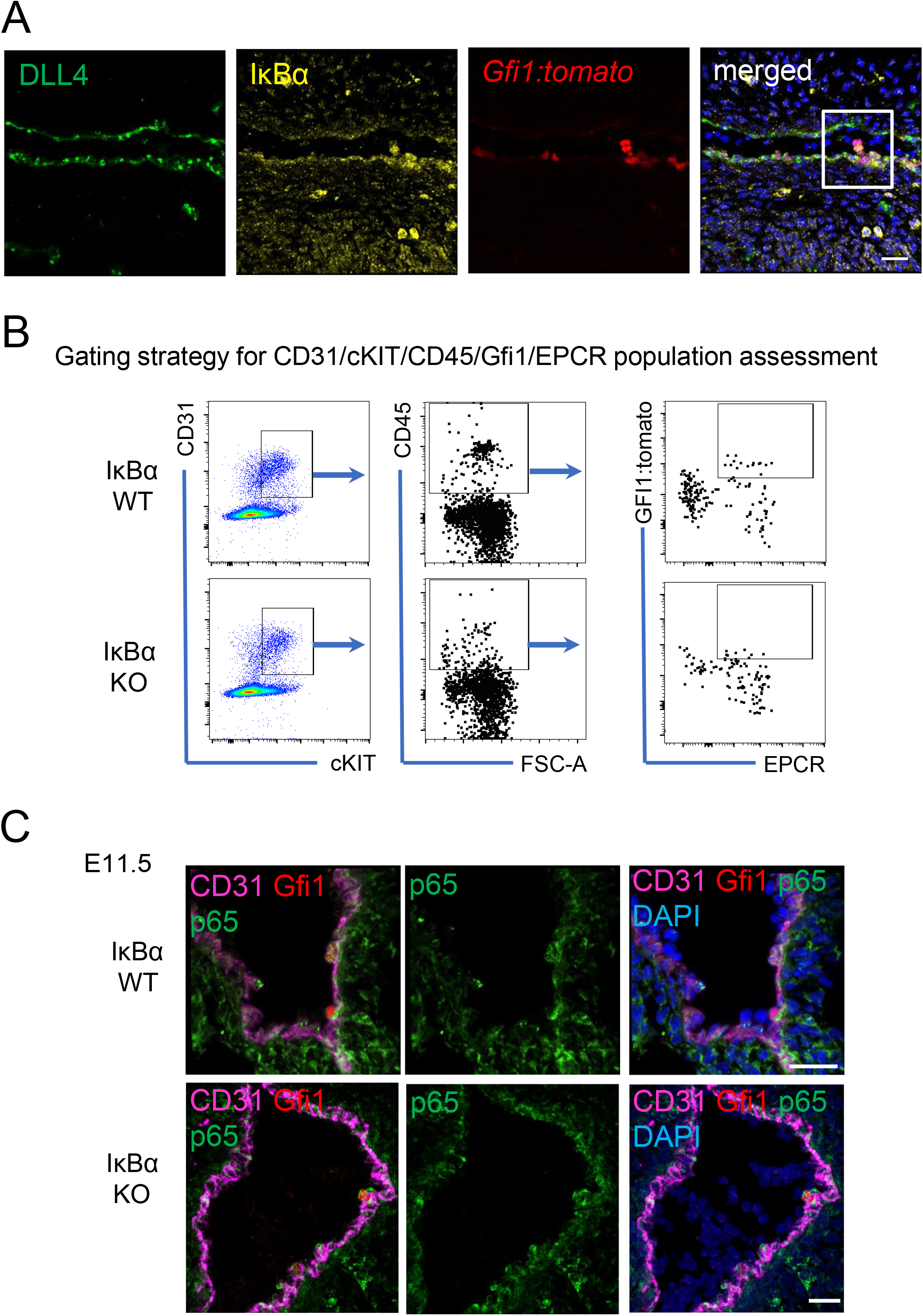
IκBα shows nuclear staining in Gfi1+ HE/IAHC but p65 staining is not altered in IκBα KO. **(A)** IHC on E11.5 (43-45s) sagittal AGM section for DLL4 (green) and IκBα (yellow) and DAPI (blue) on *Gfi1:tomato* (red) transgenic embryos. Scale bar: 60um. **(B)** Representative FACS plots for CD31/cKIT/CD45/Gfi1/EPCR+ staining in E11.5 AGMs of IκBα WT/Het and KO. **(C)** IHC on E11.5 (43-45s) AGM section for CD31 (magenta) and p65 (green) and DAPI (blue) on *Gfi1:tomato* (red) transgenic embryos. Scale bar: 40um (top panel), 60um (bottom panel).

**Suppl Figure S3:**
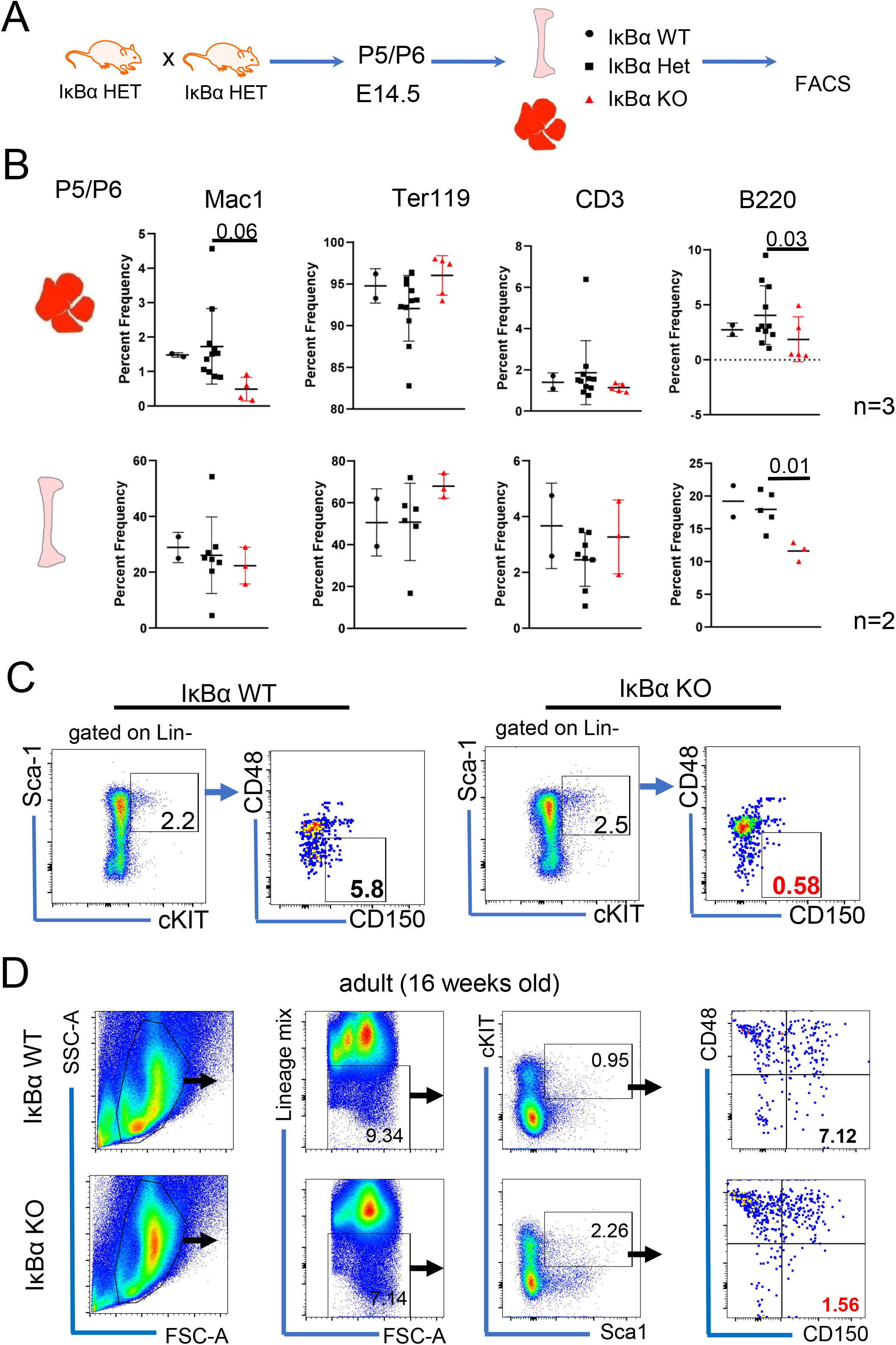
Blood lineage profiling of newborn IκBα KO and E14.5 foetal liver. **(A)** Scheme of mating’s to obtain IkBα WT, Het and KO hematopoietic tissues. **(B)** Bar chart with individual values for the frequency of Ter119 (erythoid cells) B220 (B-cells), CD4/8 (T cells) and Gr1/Mac1 (myeloid cells) in foetal liver (top) and bone marrow (bottom) of newborn (P5/6) IκBα WT, Het and KO, n=26 pups in 3 independent experiments. n= 4 WT and 6 KO for foetal liver and 9 pups in 2 independent experiments for bone marrow. P-value determined with two tailed t-test. **(C)** Representative FACS plots gating strategy for LSK (lin-Sca+cKIT+) and LT-HSC (LSKCD48-CD150+) of IκBα WT, Het and KO bone marrow and (foetal) liver. **(D)** Representative FACS plots for the frequency of LSK and LT-HSC in one adult (3 months old) bone marrow of IkBα WT and KO.

**Suppl Figure S4:**
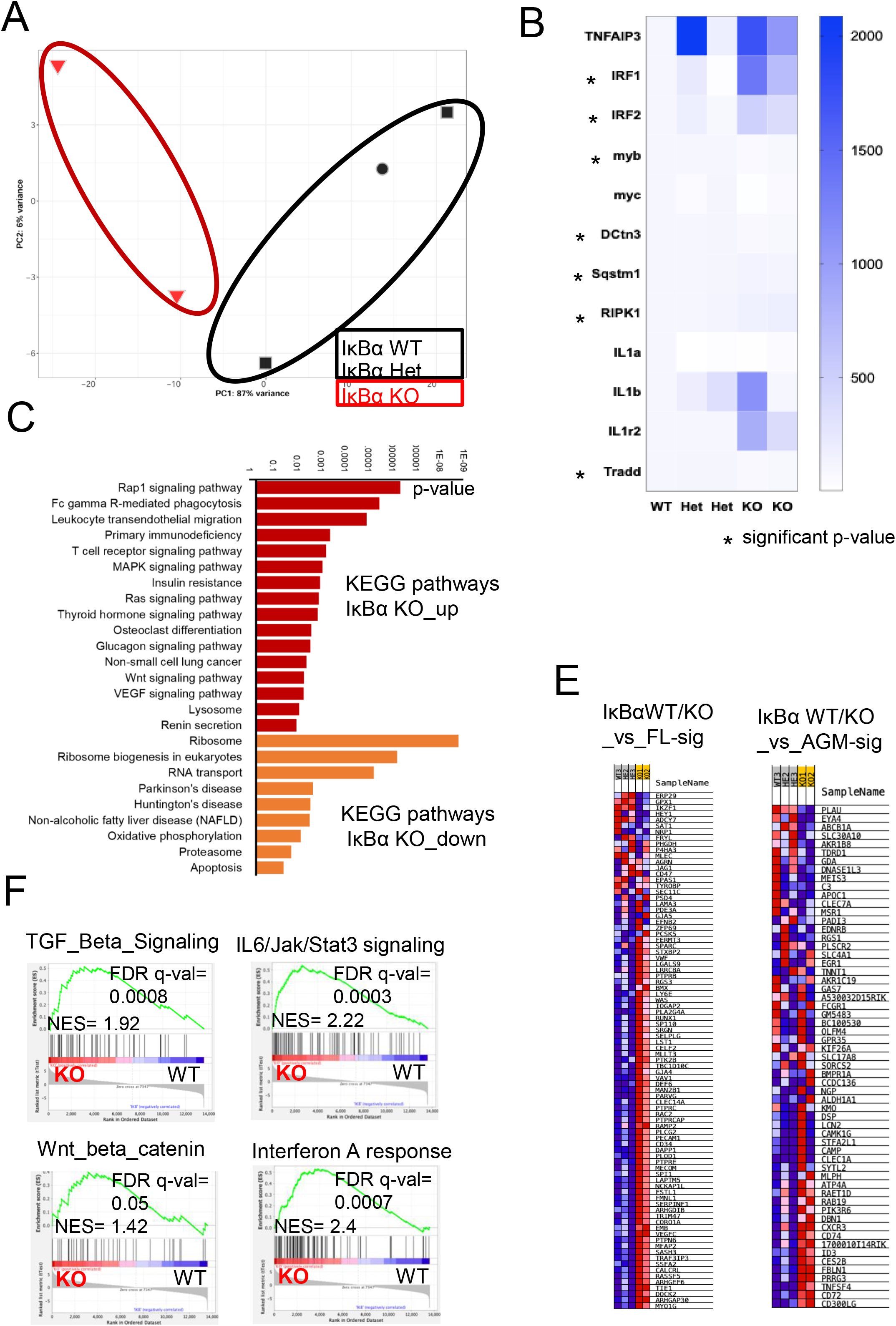
Molecular characteristics of E14.5 IκBα KO LT-HSCs. **(A)** Principal Component clustering of the LT-HSC RNA sequencing samples from 1 WT, 2 Het and 2 KO for IκBα. **(B)** heatmap with the reads counts of NFκβ related genes in LT-HSCs of IκBα WT/Het and KO. Star depicts significant p-value as determined with two tailed t-test. **(C)** Bar chart showing the KEGG pathways, and their p values associated with genes expressed at a lower/higher levels (</>2^log2^) in IκBα KO compared to IκBα WT. **(D)** GSEA of genes at a higher level (>2^log2^) in IκBα KO compared to IκBα WT.**(E)** Heat map correlation of genes that are upregulated in LT-HSCs at E14.5 IκBα KO and signature genes for foetal liver (E14.5) LT-HSCs.

**Suppl Figure S5:**
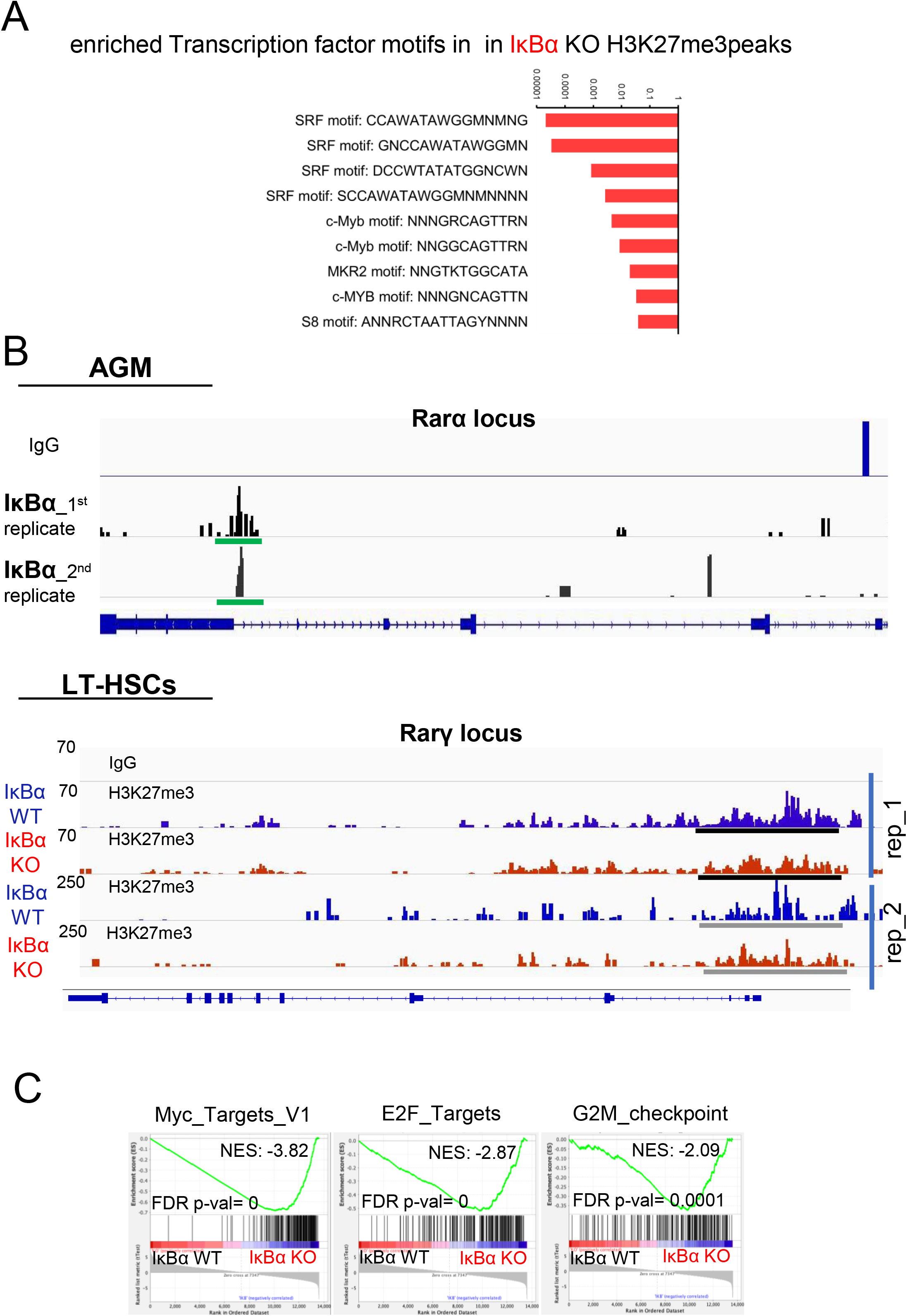
IκBα binds to retinoic acid receptors rarα and rarγ. (**A**) Bar chart showing the transcription factor motifs and their p values associated with H3K27me3 peaks that are unique to IκBα KO. (**B**) Snapshots from igv browser depicting enrichments obtained from 2 independent cut and tag assays with E11.5 CD31+ cells with the IκBα antibodies at the rarα locus (green bars indicate peaks), and igv browser snapshot from the rarγ locus in E14.5 FL LT-HSC. Black bars indicate enrichment of HH3K27me3 in IκBα WT ot KO. (**C**) Selected GSEA plots of genes expressed at lower levels in IκBα KO LT-HSC.

**Suppl Figure S6:**
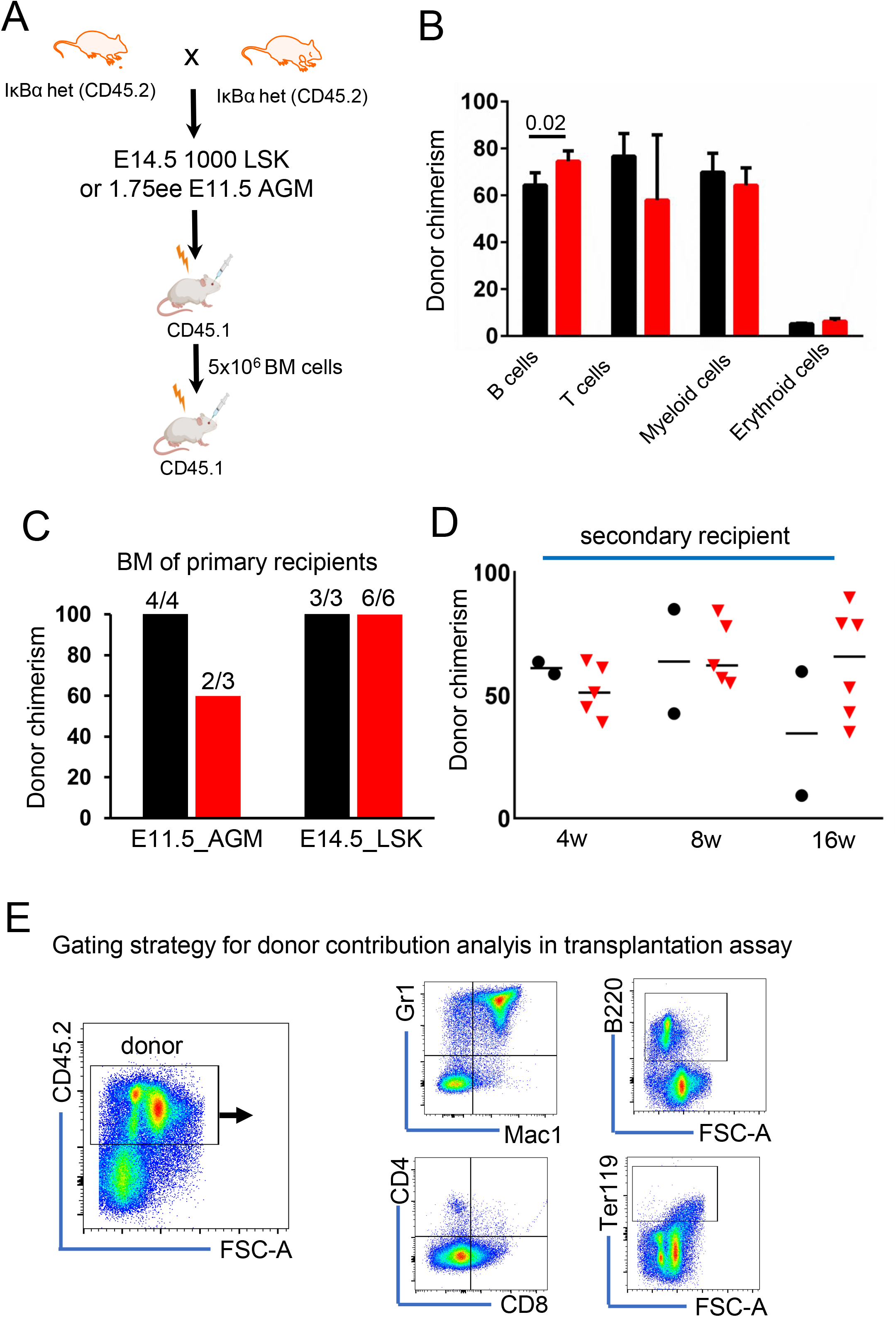
IκBα KO HSC are activated with a delay. (**A**) Scheme of transplantation assay to test hematopoietic function of IκBα WT and KO HSCs. (**B**) Donor chimerism in the bone marrow of primary recipients n=16 foetal livers in 2 independent experiment. (**C**) Bar chart of donor chimerism in secondary recipients in 4 week intervals after receiving 5x10^6^ IκBα WT or KO (CD45.2) of primary recipients of 1000 IκBα WT or KO LSKs. (**D**) Bar chart of donor chimerism in the different blood lineages at 16 weeks after receiving 1000 IκBα WT or KO LSKs (CD45.2) together with 500.000 BM cells as competitors (CD45.1). Spleen was assessed for B cells (B220), PB for Ter119 (erythoid cells), PB for CD4/8 (T cells) and BM for Gr1/Mac1 myeloid cells n= 4 WT and 6 KO in one experiment. P-value determined with two tailed t-test. **(C)** Representative FACS gating plots for the analysis of donor chimerism to the different blood lineages in transplantation assays.

## Materials and Methods

### Mouse lines and animal work

The *Gfi1:tomato* (Thambyrajah et al., 2016) and IκBα targeted knock out mouse line (Nfkbia ^tm1Bal^, Jackson Laboratories, stock #002850) were used in this study. For time matings, *Gfi1:tomato*^*tomato*^ or IκBα het and females were mated to *Gfi1*^*tomato*^ or IκBα het males. Vaginal plug detection was considered as day 0.5. The resulting embryos were genotyped and used for IHC, FACS based sorts and transplantation assay. Animals were kept under pathogen-free conditions, and all procedures were approved by the Animal Care Committee of the Parc de Recerca Biomedica de Barcelona, license number 9309 approved by the Generalitat de Catalunya.

### E11.5 AGM transplantation experiments

AGMs of E11.5 embryos (CD45.1/2) were dissected in PBS with 7% fetal calf serum (FBS) and penicillin/streptomycin (100 U/mL). Single cell suspensions were generated by incubating the tissues for 20-30 minutes in 500 ul of 1mg/ml of Collagenase/Dispase (Roche cat# 10269638001) before mechanical dissociation with a syringe and needle. The resulting single cell suspension was used for transplantation assays. Briefly, Charles River NSG mice (CD45.2) were irradiation at 4 Gy and 1.75 embryonic equivalent AGM was injected retro-orbitally into each recipient, along with 20.000 bone marrow cells from a competitor (CD45.1). Peripheral blood (PB) donor chimerism was analyzed by FACS at 4, 8, 12, and 16 weeks. Lineage analysis on BM-derived cells was performed at 16 weeks post-transplant by flow cytometry with specific antibodies. Please see Suppl table S4 for a list of antibodies. The samples were analyzed on a Fortessa or LSRII instrument and FlowJo v10. Statistical significance was determined with GraphPad prism 8.

### Foetal liver and bone marrow transplantation experiments

Charles River C57BL/6 (CD45.1) was used as recipients after two rounds of irradiation at 4 Gy (8 Gy total). FACS isolated LSK cells (1000), or purified ST-HSC (LSKCD48+CD150) and LT-HSC (LSKCD48-CD150+) from IκBα WT, Het or KO (CD45.2) fetal liver were retro-orbitally injected with 500.000 lineage depleted bone marrow cells as competitor (CD45.1) foe primary transplantation assays. In secondary recipients (CD45.1), the bone marrow of the primary recipients (CD45.2) was extracted at 16 weeks and a total of 5 million cells were transplanted into each secondary recipient after a total of 8 Gy of irradiation. Peripheral blood (PB) donor chimerism was analyzed by FACS at 4, 8, 12, and 16 weeks. Lineage analysis on BM-derived cells was performed at 16 weeks post-transplant by flow cytometry with specific antibodies. Please see Suppl table S4 for a list of antibodies. The samples were analyzed on a Fortessa or LSRII instrument and FlowJo v10. Statistical significance was determined with GraphPad prism 8.

### Genotyping PCR

Small pieces of embryonic tissue or yolk sac were dissected off the embryo and placed in PCR tube containing 30μl of PBS. The tissue pieces were boiled for 8 minutes at 98°C for denaturation. The tissues were digested with Proteinase K (50 μg/ml) for 30 minutes at 55°C, and the enzyme deactivated by boiling the samples for a further 10 minutes at 95°C. 1μl of the samples was used as a template for the PCR.

### AGM dissection, single cell suspension

AGMs of E10 - E11.5 embryos were dissected in PBS with 7% fetal calf serum (FBS) and penicillin/streptomycin (100 U/mL). Single cell suspensions were generated by incubating the tissues for 20-30 minutes in 500 ul of 1mg/ml of Collagenase/Dispase (Roche cat# 10269638001) before mechanical dissociation with a syringe and needle. The resulting single cell suspension was used for antibody staining (see Suppl table T4 for list of antibodies). samples were analysed on a Fortessa instrument and further plotted using FlowJo V10.

### Lineage committed cell depletion from foetal liver and bone marrow

Mouse bone marrow cells were obtained by extracting the femur and tibia bones and subsequent crushing of the bones in cold PBS with 0.5% FBS Buffer using mortar and pestle. Cells were then labeled with a cocktail of antibodies against lineage markers (BD Bioscience, cat# 559971) before conjugation to anti Biotin microBeads (Miltenyi Biotech, cat #130-090-485). Labeled cells were depleted using magnetic columns (Miltenyi Biotech), leaving behind isolated unlabeled Lin– cells immediately ready for downstream experiments. For FL, the tissue was harvested from E14.5 embryos and the dissected liver was crushed with the end of a 1ml syringe through a 40um cell strainer into IMDM +10% FBS. For FL stage, CD11b was not included in the lineage depletion mix.

### Flow Cytometry and cell sorting

Sorts of bone marrow cells were performed on FACSAriaII or BD Influx (BD Biosciences) instrument after magnetic depletion of lineage primed celled (Miltenyi Biotec) and staining with the appropriate antibodies in PBS with 10% FBS (see Suppl table T5 for list of antibodies).

### Single Cell LT-HSC culture ex vivo

The foetal liver was harvested from E14.5 embryos and depleted from linage primed progenitors as described above. Subsequently, samples were stained for LSKCD48CD150 and sorted as single cells (see above) into 96 well plates containing complete Stem Cell Medium (StemSpan SFM, LifeTechnologies containing 50 ng/ml IL-3, 50 ng/ml IL-6, 2 ng/ml FGF2, 50 ng/ml SCF, 25 ng/ml TPO, 30 ng/ml Flt3-Ligand (all Preprotech), 100 u/ml Penicillin/Streptomycin, 2 mM L-Glutamine). Cells were cultured in 96-well ultralow attachment plates for 10 days and colonies were scored under a light microscope.

### Ex vivo treatment of LT-HSC with Ro-41

The foetal liver was harvested from E14.5 embryos and depleted from linage primed progenitors as described above and were treated with either Ro 41-5253 (Sigma-Aldrich) (10 μM), or the respective amount of DMSO.

### Cell cycle analysis

For cell cycle analysis, samples were stained for LSKCD48CD150 first and further stained for the intracellular anti-Ki-67 Alexa Fluor® 647 (Clone B56, BD Pharmingen) and DAPI using the FIX & PERM™ Cell Permeabilization Kit from Thermo Fisher (cat# GAS003). The samples were run on a Fortessa Instrument from BD Bioscience and analyzed with FlowJO V10. Statistical significance was determined with GraphPad prism 8.

### BrdU labelling/staining

2mg pf BrdU was injected in a total volume of 400ul into pregnant females 2 hours before sacrifice. The foetal samples were processed and stained according to manufacturer’s recommendations (APC BrDU Kit, BD Bioscience, cat # 552598). The samples were run on a Fortessa Instrument from BD Bioscience and analyzed with FlowJO V10. Statistical significance was determined with GraphPad prism 8.

### Immuno-Histochemistry

E10.5 embryos were fixed in 2% Paraformaldehyde (Thermo Fisher) for 12 minutes, before they were soaked in 30% sucrose overnight and mounted in OCT compound. 10-12 μm sections were prepared using a cryostat. The sections were streptavidin/biotin blocked if a biotin antibody was used followed by serum blocking (PBS with 10% FCS, 0.05% Tween20) for 1 hour before the sections were incubated with primary antibodies at 4°C overnight in blocking buffer. Sections were washed three times in PBST for 15 minutes each and then incubated with fluorochrome-conjugated secondary antibody at room temperature for 1 hour. Sections were further washed three times in PBST and mounted using Prolong Gold anti-fade medium with DAPI (Life Technologies). Images were taken using a SPE (Leica) and processed using Imaris v4.8.

### Cut and Tag assay

For Cut and Tag assays, either CD31+ (25.000-30.000 cells) from E11.5 AGMs or LT-HSCs from E14.5 foetal livers (3.000-4.000 cells of IκBα WT or KO) were FACS sorted and used for cut and tag assay of IκBα and H3K27me3, respectively. We followed the Bench top CUT&Tag V.3 (https://www.protocols.io/view/bench-top-cut-amp-tag-bcuhiwt6), an updated version from Kaya-Okur et al, 2019 with their recommended antibody for H3K27me3 (for details, please see Suppl table T5) and loaded Tn5 from epicypher. After the assay, the samples were quantified and validated using a bio analyzer and subsequently sequenced as a 2x50 bp run on a HiSeq (Illumina).

### Cut and Tag assay peak calling

Raw single-end 50-bp sequences were filtered by their quality (Q>30) and length (length > 20-bp). Filtered sequences were aligned against the reference genome (mm10 release) with Bowtie2.Basecall files generated from HiSeq sequencing run were converted to FASTQ format with Illumina’s bcl2fastq. Alignment was performed by bowtie2 (version 2.2.1) to mouse reference genome mm10 with default parameters. Generated SAM files from bowtie2 alignment were converted to BAM files by samtools v0.1.19. Parameters for samtools SAM to BAM conversion: -bS -q 10 -F 260. Peaks were called using the program macs2 (version 2.1.0.20150420) with the settings “-q 0.0085”. Full pipeline repo available at: https://github.com/mproffitt/BioWorkflow. Overlaps of gene list were performed with venny venn 2.0 online (https://bioinfogp.cnb.csic.es/tools/venny/index2.0.2.html) and KEGG pathways associations were determined with DAVID pathway analysis tool (https://david.ncifcrf.gov/tools.jsp). Peak visualization was done with Integrative Genomics Viewer (IGV). Cut and tag-sequencing data are deposited at the GEO database with accession number GSE188525.

### RNA-sequencing data analysis

In all experiments, we extracted total RNA from three mice per genotype using the mRNA ultra low input protocol for library prepreparation (Quiagen, cat# 180492). The RNA concentration and integrity were determined using Agilent Bioanalyzer [Agilent Technologies]. Libraries were prepared at the Genomics Unit of PRBB (Barcelona, Spain) using standard protocols, and cDNA was sequenced using Illumina® HiSeq platform, obtaining ∼ 25–30 million 50-bp single-end reads per sample. Adapter sequences were trimmed with Trim Galore. Sequences were filtered by quality (*Q* > 30) and length (> 20 bp). Filtered reads were mapped against the latest release of the mouse reference genome (mm10) using default parameters of TopHat (v.2.1.1), and expressed transcripts were then assembled. High-quality alignments were fed to HTSeq (v.0.9.1) to estimate the normalized counts of each expressed gene. Differentially expressed genes between different conditions were explored using DESeq2 R package (v.1.20.0) <u>36</u>. Plots were done in R. RNA-sequencing data are deposited at the GEO database with accession number GSE188525.

